# Rapid stem cell spreading induced by high affinity α_5_β_1_ integrin-selective bicyclic RGD peptide in biomimetic hydrogels

**DOI:** 10.1101/2022.02.01.478177

**Authors:** Kaizheng Liu, Johannes Vandaele, Dominik Bernhagen, Merijn van Erp, Egbert Oosterwijk, Peter Timmerman, Susana Rocha, Paul H. J. Kouwer

**Affiliations:** Radboud University, Institute for Molecules and Materials, Heyendaalseweg 135, 6525 AJ Nijmegen, The Netherlands; KU Leuven, Department of Chemistry, Molecular Imaging and Photonics, B-3001 Leuven, Belgium; Pepscan Therapeutics, Zuidersluisweg 2, 8243 RC Lelystad, The Netherlands; Radboud University Medical Centre, Radboud Institute for Molecular Life Sciences, Microscopic Imaging Centre, Nijmegen, The Netherlands; Radboud University Medical Centre, Radboud Institute for Molecular Life Sciences, Department of Urology, Geert Grooteplein 26-28, PO Box 9101, 6500 HB Nijmegen, The Netherlands

## Abstract

Cell-matrix interactions form a crucial parameter for the design of a synthetic extracellular matrix (ECM), as they ultimately dictate cell fate and functions. Universally, synthetic biomaterials are conjugated with a linear or cyclic Arg-Gly-Asp (RGD) peptide to establish a direct link of the ECM with the cell. These peptides, however, present low binding affinities and lack selectivity towards integrin subtypes presented on the cell membrane. Here, a highly biomimetic synthetic ECM based on polyisocyanides (PIC) that has been decorated with bicyclic peptides that show a high affinity towards specific integrin subtypes is presented. 3D cell studies show that human adipose-derived stem cells (hASCs) in matrices carrying the optimized bicyclic α_5_β_1_ -integrin binder, spread within 24 hours, which is much faster than in other PIC gels, including the default RGD-decorated gel, but also much faster than in the positive Matrigel control. YAP/TAZ staining shows that the rapid morphological change in the 3D microenvironments is YAP independent. The data highlights that the design of synthetic matrices with appropriate, optimized guiding signals is key to guide cells towards a predetermined outcome.

## Introduction

Human tissue is a complex system consisting of cells and the extracellular matrix (ECM). The latter supports the structural integrity of the tissue and has profound effects on the embedded cells. For instance, the ECM supports cell adhesion, regulates growth factor presentation to mediate cell functions and is involved in the activation of many cell signaling pathways^1^. Cells sense their microenvironment and show a corresponding biological response^2^.

Integrins form the physical connection between a cell and its extracellular matrix. They are allosteric transmembrane proteins that mediate bi-directional interactions between cells and their (micro)environment and, as such, they play a central role in communication^3,4^. On the intracellular side, integrins interact with a range of adhesion-supporting proteins and signaling factors to form force-responsive adhesion complexes. These complexes bind the actomyosin cytoskeleton, regulating downstream signaling pathways that allow the cell to react to physical cues. Reciprocally, on the extracellular side where the integrins bind to the ECM, cell contraction leads to a change in the architecture and mechanical properties of the cellular microenvironment.

Functional integrins consist of two non-covalently bound transmembrane subunits, named α and β. On human cells, 24 heterodimers have been observed experimentally, which are composed of combinations of 18 α and 8 β subunits^5,6^. Cells have phenotype-dependent presentation of integrins and the integrin expression may change in physiological or pathological processes, for instance, during stem cell differentiation^7^ or in cancer^8^.

Interestingly, selectivity is not always high for ECM-integrin interactions, e.g., collagen and laminin, two major ECM proteins, both bind α_1_β_1_ and α_2_β_1_ integrins. Laminin additionally binds with α_5_β_1_ integrins, which in turn, also interacts with fibronectin. In 1984, a tripeptide sequence Arg-Gly-Asp (RGD) was identified as a principle integrin-binding domain^9^. This peptide, subsequently, has become the golden standard in research involving cell-adhesion^10^, particularly in synthetic microenvironments. The advantages of this classic sequence include its stability during sample preparation and processing, its minor immune response and the availability of simple and economical conjugation strategies, for instance, *via* click chemistry^11^. Together, they make RGD and its derivatives the primary choices to induce a cell-adhesion in a non-biofunctional material. The simple RGD peptide also has some clear drawbacks. In the absence of a secondary structure of the RGD peptide, cell anchoring through the integrin is not comparable to the full-length proteinprotein^12,13^. Furthermore, RGD binds generically to multiple integrin subtypes and cannot be used when a specific heterodimer needs to be targeted, which limits the possibility of activation of specific desired intracellular pathways.

To pursue a stronger and more selective binding towards integrins, cyclic peptides have been developed^14^. In 2D *in vitro* settings, these peptides have been shown to promote cell adhesion to gel substrates^15,16^ and cell membrane binding^14^. In this work, we introduce combinatorically optimized cyclic peptides in a highly biomimetic, but fully synthetic hydrogel based on polyisocyanides to study cell behavior in a controlled *in vivo*-like microenvironment. Our first results show the dramatic effects that binding optimization offers but, ultimately, we present a new method that allows one to start targeting specific integrins to guide desired cell behavior such as migration, stem cell differentiation, and angiogenesis^17^.

## Results and Discussion

As the synthetic biomimetic matrix, we employed polyisocyanide (PIC) gels^18^. This polymer-based fibrous hydrogel is developing into an ideal synthetic three-dimensional artificial ECM^19-21^. PIC gels combine an architecture and (nonlinear) mechanics that are common to hydrogels of natural ECM polymers and possess excellent biocompatibility^22^ with high customizability in mechanical properties and biofunctionalization.

We exploited the straightforward customization strategies to introduce 9 different adhesive peptides (**Table 1**), including the routinely used linear GRGDS (**P1**), the cyclic fKRGD (**P2**), the knottin peptide (**P3**, optimized for strong integrin binding) and a series of recently reported constrained, bicyclic peptides^14,23^ that have been optimized for strong and selective integrin binding^24^ (**P4-P9**). For conjugation, we employed the highly efficient strain-promoted azide-alkyne cycloaddition (SPAAC) reaction^25,26^ between the azide (N3)-functionalized PIC polymers and the bicyclononyne-terminated peptides^16^ using earlier developed protocols^19^. To drive the conjugation reaction to completion, an excess of N3 groups is used^27^. The design of our 3D cell culture model system is displayed in **Figure 1**. The final peptide concentration after cell encapsulation is 63 μM, which is considered low, compared to other synthetic matrices^28-31^.

**Table 1.**
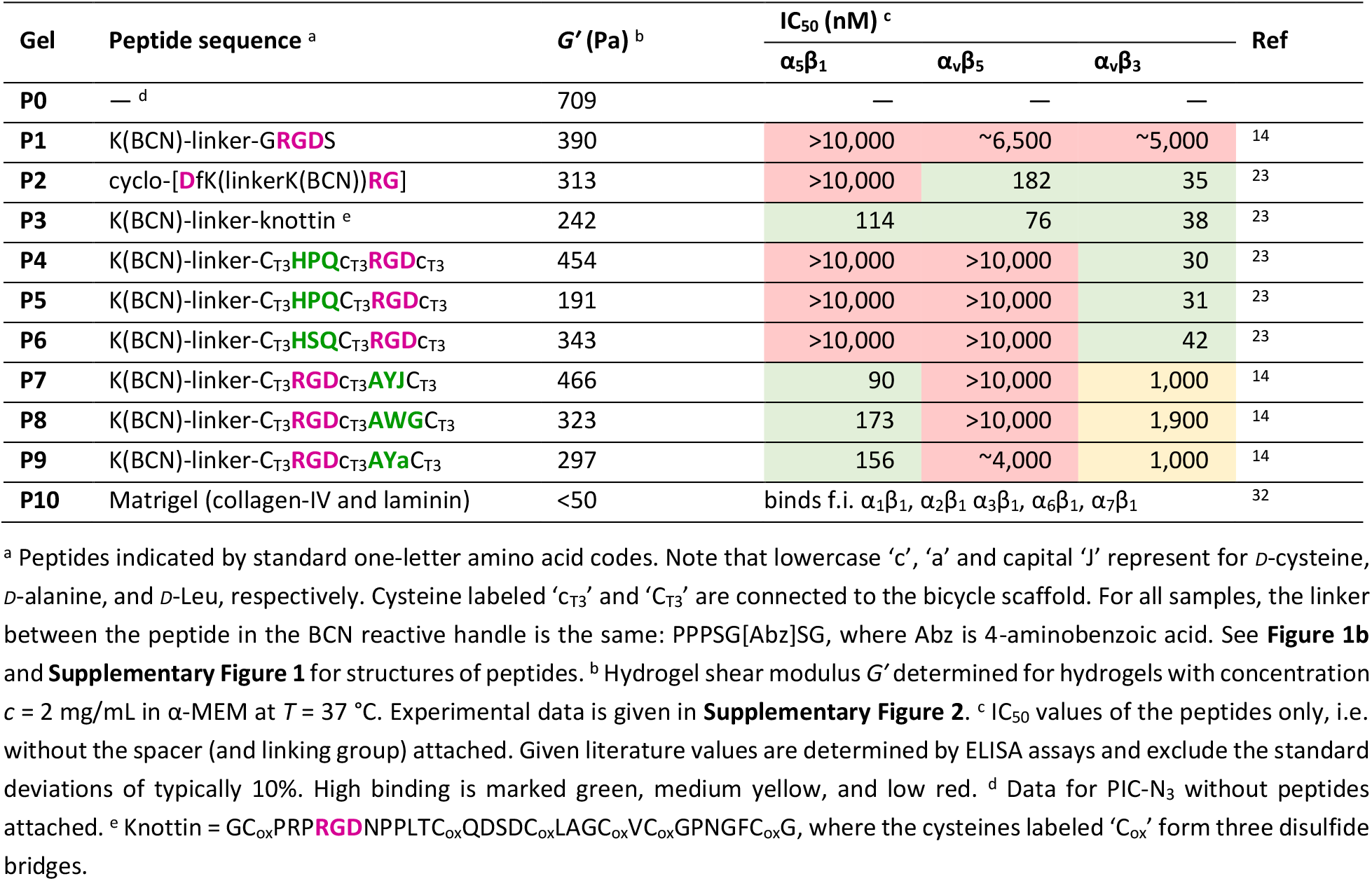
Overview of hydrogels **P1**-**P10** prepared after conjugation of PIC-N_3_ with the appropriate peptide sequence.

**Figure 1.**
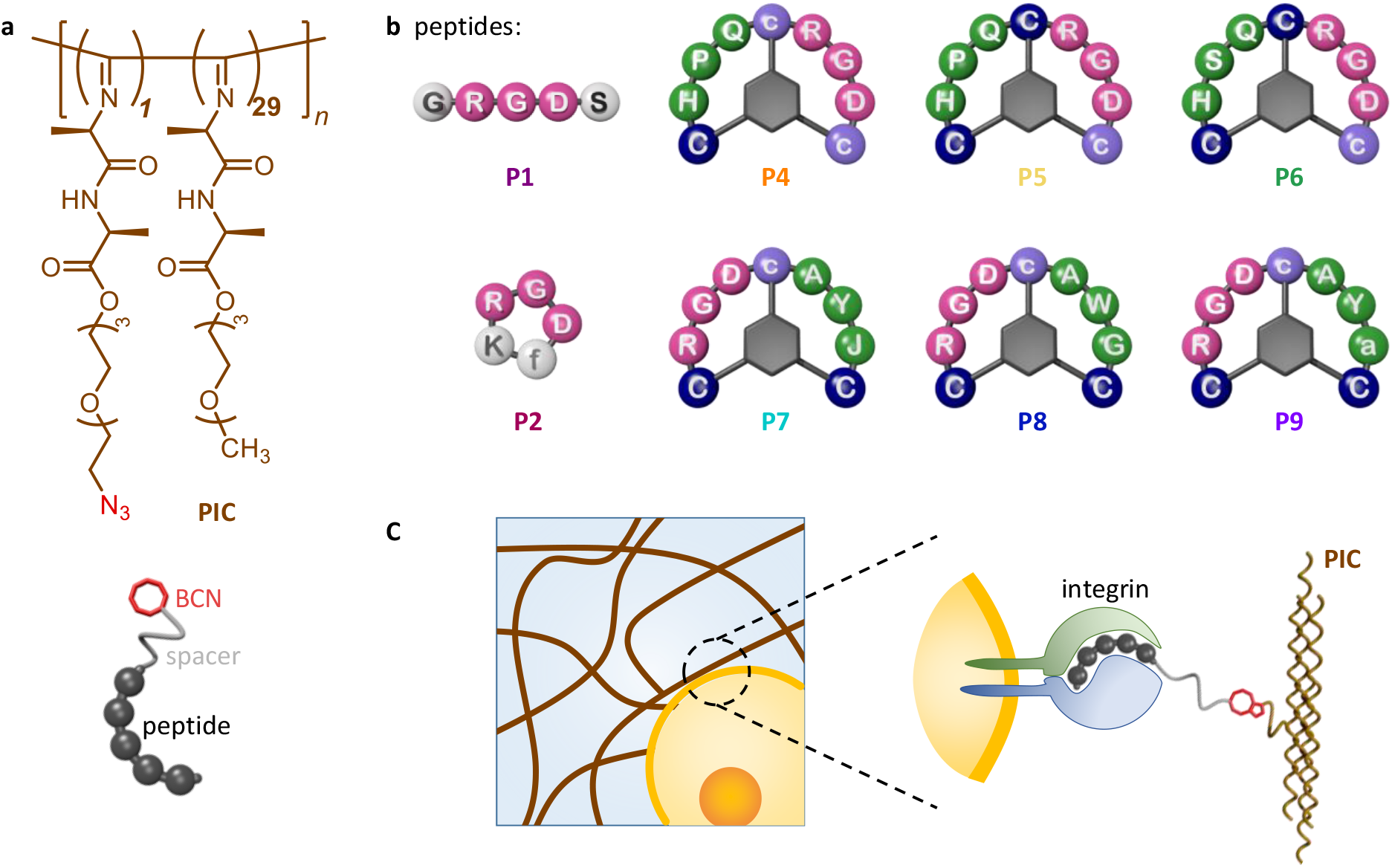
Design of the cell culture materials. **a**, Structure of the PIC polymer with an azide (N_3_) that serves as a reactive handle for bio-orthogonal click-chemistry and, schematically, the linker-peptide construct with reactive BCN (red) end: K(BCN)PPPSG[Abz]SG-peptide, where Abz is 4-aminobenzoic acid. For PIC, the ratio between azide and methoxy-terminated monomers is set to 1:29 (*x* = 0.033), but only ∿1% of the monomer later is substituted with a peptide. **b**, Structure of linear (**P1**), monocyclic (**P2**) and bicyclic peptides (**P4**–**P9**) using single letter amino acid codes. Note that lowercase ‘c’, ‘f’ and capital ‘J’ represent *D*-cysteine, *D*-phenylalanine and *D*-leucine, respectively. **P3** is not presented in this figure due to its complex structure. **c**, For stem cell studies, we generated a synthetic fibrous microenvironment with a single cell adhesion peptide clicked onto the polymer backbone.

PIC hydrogels were prepared by dissolving the (peptide-functionalized) polymer in cold cell culture medium and heating the solution beyond the gelation temperature *T*gel ≈ 15 °C. For all hydrogels, the low peptide concentrations do not influence the gelation temperature of the polymer. At 2 mg mL^−1^ and 37 °C, all PIC gels form relatively soft hydrogels with storage moduli between 200–500 Pa, depending on the peptide attached. Note that the PIC polymer without peptides forms a slightly stiffer gel (**P0**).

Earlier reports quantified the binding of the peptides towards different target integrins: α5β1, αvβ3 and αvβ5 (**Table 1**). The routinely applied linear GRGDS binds weakly to all integrins; while the frequent applied cyclic RGD (in **P2**) has a higher binding affinity, but only towards the αvβ3 and αvβ5 heterodimers. The engineered knottin RGD, however, strongly interacts with all three integrin dimers (α5β1, αvβ3 and αvβ5). Of the two groups of optimized bicyclic peptides, the first group selectively binds αvβ3 integrins (**P4**–**P6**) while the second is selective for α5β1 (in **P7**–**P9**). Matrigel (**P10**) was used as the ‘golden standard’ reference material in the 3D cell culture field.

We tested the role of the integrins in cell culture with human adipose-derived stem cells (hASCs). Earlier work has shown that somatic stem cells are highly sensitive to their (mechanical) microenvironment^33^, to which they anchor through their integrins. In the case of hASCs, it has been observed that α5β1 is highly expressed during the undifferentiated state^7^. We encapsulated hASCs in (200,000 cells mL^−1^) in every gel and cell viability (LIVE/DEAD staining) measurements after 72 h indicate that all gels are fully biocompatible (**Figure S3**).

The influence of the different integrin binding peptides in cellular behavior was evaluated by monitoring cell morphological responses (spreading and migration) in the first 24 hours after encapsulation. We focused on the initial time period to minimize the contribution of the extracellular matrix deposited by the stem cells. Analysis of bright field images (**Figure S4**) shows that 24 hours after encapsulation, the stem cells are still primarily spherical in all matrices, apart from **P8**, where a network of spread cells was imaged. Based on the IC50 values, gels with **P8** are expected to have a strong interaction with the α5β1 integrin, which is expressed by the stem cells. While **P7** and **P9** are decorated with peptides that bind similarly strong, the encapsulated cells in the corresponding gels remain mostly spherical. Similarly, in gels with the generic strong binder **P3**, no morphological changes are observed. We underline that integrin binding constants were determined by ELISA with the peptides only and that conjugation of the peptides to a polymer backbone may induce a different functionality as is sensed by the cells.

Based on this data, we selected 5 culture matrices for more quantitative analysis of the role of the peptide on cell behavior: **P0** (no peptide, negative control), **P1** (default GRGDS peptide), **P4** and **P8** (bicyclic peptides with high affinity for either the αvβ3 or the α5β1 integrin receptor, resp.) and Matrigel (**P10**) as a biological reference material containing abundant collagen (affinity towards α2β1 integrins)^34^, and laminin (that binds α1β1, α2β1 α3β1, α6β1 and α7β1 integrins)^35^. Time-lapse bright field microscopy (**Supplementary Movies M1–M5**) clearly shows the remarkably fast spreading of cells in **P8**, in contrast to the other samples (**Figure 2a** shows images at 7 and 24 h; **Figure 2b**, images at 1, 3, 5 and 7 h of **P8** only). As a measure for spreading, we determined cell circularity for ∼20 representative and isolated cells. Quantitative analysis using FIJI (**Figure 2c**), indeed confirms that already after 3 h, hASCs in **P8** are significantly different from cells in any of the other matrices, including Matrigel (**Figure 2c**). Even at 24 h, the cells in the other matrices still show a rounded morphology, compared to a well-developed cellular network in **P8**. Confocal fluorescence imaging of F-actin stained samples (24 h) confirms the results observed in the bright field images (**Figure 2d**). Experiments to dilute the density of peptides in **P8** (by culturing hASCs in mixed **P8**/**P0** hydrogels) shows that 63 μM is the minimum peptide concentration to obtain extensive spreading (**Figure S5**).

**Figure 2.**
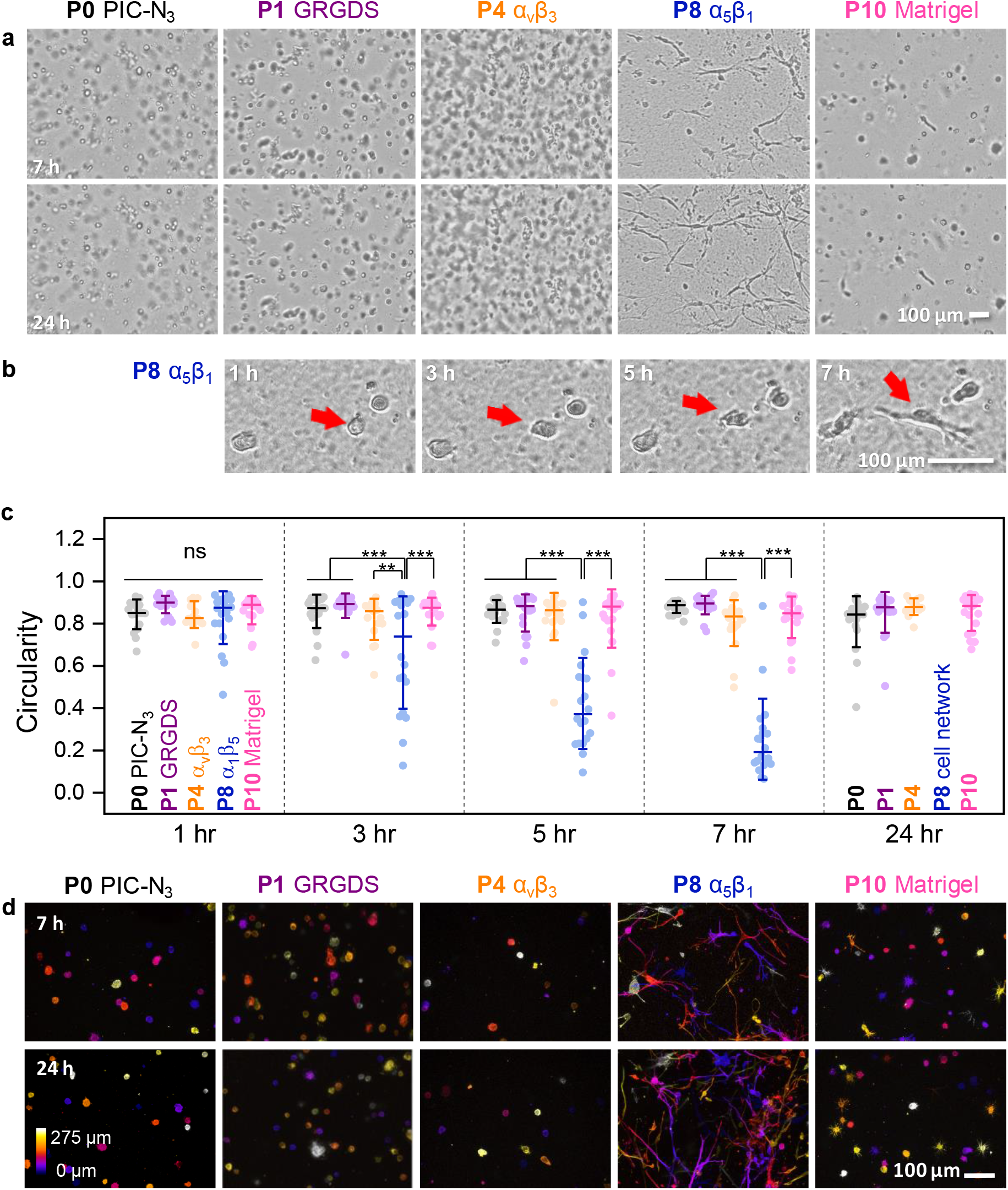
Cell spreading in gels functionalized with different peptides. **a**, Bright field images of stem cell cultures **P0, P1, P4, P8** and **P10** at 7 and 24 h after encapsulation. **b**, Representative bright field images show the morphological change of stem cells in the selective α_5_β_1_ binder gel **P8** at 1, 3, 5 and 7 h after encapsulation. **c**, Quantification of cell circularity as a measure for spreading, based on bright field images. Box plot shows data points, median and ± SD. For all cultures, ∿20 non-overlapping cells are traced for 24 h. Statistics: ** P<0.01, *** P<0.001. **d**, False color-coded (depth) confocal fluorescence microscopy projection images showing F-actin organization after 7 and 24 h after encapsulation. For all cell cultures: PIC gel concentration 2 mg mL^−1^, adhesive peptide concentration 63 μM and cell encapsulation density 200,000 cells mL^−1^.

Yes-associated protein (YAP) and the co-activator TAZ are key mechanotransduction components that translocate from the cytosol to the cell nucleus upon mechanical cues from the cellular microenvironment^36^. Given the rapid cell spreading in **P8**, we evaluated YAP activation in our samples. Interestingly, immunostaining reveals that there is no significant YAP localization in cell nuclei in either condition, implying that within 24 hours, the YAP/TAZ mechanotransduction pathway is not activated in hASCs in these 3D hydrogels (**Figure S6**). Our results contradict findings from 2D cell culture^36^ that can give rapid nuclear translocation but are in line with the other recent findings from 3D cell cultures, where no redistribution of YAP is observed^37^. Indeed, the lack of actin stress fibers, the decrease of nuclear content per cross-section and low volumes of the cells may cause the difference between the 2D and 3D microenvironments. Also, we suspect that, as a result of the short culture time, the YAP pathway may not have been activated yet. In conclusion, the data suggest that YAP/TAZ is not involved in the observed rapid spreading of stem cells in the first 24 hours.

The capacity for matrix remodeling and migration are fundamental stem cell characteristics that are necessary to carry out their function^38^. The migration of cells in a 3D microenvironment is an extremely intricate reciprocal process^39^, which also includes matrix degradation. Integrin-RGD complexes are central to mechanical sensing of the microenvironment, and at the same time, they also function as traction points that are needed for cell movement^38^. The migration observed during cell spreading in **P8** is cell-adhesion dependent but degradation independent, as the synthetic PIC gel cannot be degraded by proteases.

We emphasize that the matrix remodeling and cell spreading/migration modes in Matrigel differ from the ones in PIC gels, as reflected by the changes in cellular morphology in the two materials (**Supplementary Movies M4, M5**). In Matrigel, stem cells produce small non-directional protrusions, but no migration is observed in the first 24 h. We speculate that cells are still degrading the surrounding ECM to create extra space. In contrast to Matrigel, the PIC-based synthetic matrix allows (bio)chemical-independent remodeling, also termed physical remodeling^40^ of ECM. The bicyclic peptides present in **P8** allow the cell to induce fast physical remodeling of the matrix by applying contractile forces. We hypothesize that in **P8** the bicyclic peptide may contribute to the formation of filopodia^41^, as α5β1 is crucial for stem cell migration^42,43^. The clustering of peptides, caused by the bundling of PIC polymers into fibers during gelation, may also facilitate cell adhesion and further morphological responses^17^. Furthermore, the plasticity of PIC hydrogel enables protease-free cell spreading and even migration in a short time span^40^.

## Conclusions

Collectively, our results show that this novel combination of a synthetic network with architecture and mechanical characteristics of naturally occurring biomaterials, together with strongly-binding cyclic peptides results in an artificial ECM with properties that are well beyond what is possible with biological materials. In one gel, we observe tremendous acceleration in cell spreading, which, to our knowledge, has not been observed in any synthetic or biological matrix. Additionally, the results illustrate the importance to target specific integrin subunits using cell-adhesion peptides when designing biomaterials. Although in this work, we only vary the cell-adhesive peptide, the conjugation approach allows for the introduction of a broad variety of active biomolecules. Using this strategy, such synthetic ECMs can be modified to achieve specific properties, which opens the possibility to study cells in uniquely predefined 3D environments. For instance, stem cell migration, matrix remodeling, focal adhesion, and mechanotransduction pathways may be modeled. From a longer-term perspective, our system has the potential to be a powerful tool to manipulate stem cell behavior and be used for more tissue engineering purposes that target specific cell types.

## Materials and methods

### Polymer synthesis

Polyisocyanides were synthesized using an established protocol as reported previously^19^. In short, the isocyanide monomers (azide-appended monomer and non-functional monomer in ratio 1:29) were dissolved in freshly distilled toluene. A catalyst solution Ni(ClO_4_)_2_·6H_2_O (0.1 mg mL^−1^) in freshly distilled toluene/absolute ethanol 9:1) was added such that the total concentration monomer to Ni^2+^ ratio equaled 5000:1. Then toluene was added to adjust the final isocyanide concentration to 50 mg mL^−1^. The polymerization mixture was stirred at room temperature, and the progress of the reaction was followed by IR-ATR (disappearance of the characteristic isocyanide absorption at 2140 cm^−1^). Once the isocyanide was consumed (48 h), the polymer was precipitated in diisopropyl ether under vigorous stirring and collected by centrifugation. The polymer was dissolved in dichloromethane, precipitated for another two rounds, and air-dried to yield the polymers as off-white solids. The molecular weight of the polymer was determined by viscometry (from dilute solutions in acetonitrile 0.1-0.6 mg mL^−1^) using the empirical Mark–Houwink equation [*η*] = *KM*_v_^*a*^, where [*η*] is the experimentally determined intrinsic viscosity, *M*_v_ is the viscosity-based molecular weight. We use values of the Mark–Houwink constants *K* and *a* that were previously determined for other polyisocyanides: *K* = 1.4 × 10^−9^ and *a* = 1.75. For the polymer used in this study, we found *M*_v_ = 536 kg mol^−1^.

### Bioconjugation

The azide-appended polymer was dissolved in acetonitrile (2.5 mg mL^−1^), and the appropriate amount of the BCN–peptide (PEPSCAN, 6 mg mL^−1^ solution in DMSO, 0.33 eq. with respect to N3 groups) was added. The solution was stirred for 24 h at room temperature and the polymer–peptide conjugates were precipitated in diisopropyl ether, then collected by centrifugation and air-dried for 24 h.

### Rheology

The mechanical analysis of the gels was performed according to a previously reported protocol^19^. Briefly, a stress-controlled rheometer (Discovery HR-1 or HR-2, TA Instruments) with an aluminum or steel parallel plate geometry was used (diameter = 40 mm, gap = 500 μm). All samples were loaded onto the rheometer plate in the liquid state at *T* = 5 °C followed by a temperature ramp to *T* = 37 °C at a rate of 1.0 °C min ^−1^. The moduli were measured in the linear viscoelastic regime at an amplitude of *γ* = 0.02 or 0.04 and a frequency of *ω* = 1.0 Hz. The sample was allowed to equilibrate at 37 °C before the nonlinear measurements. Here, the gel was subjected to a constant prestress of *σ*_0_ = 0.5 to 200 Pa, and the differential modulus *K′* was probed with a small superposed oscillatory stress at frequencies of *ω* = 10 to 0.1 Hz (reported data at *ω* = 1 Hz). The oscillatory stress was at least 10 times smaller than the applied prestress.

### Cell culture and encapsulation

Human adipose-derived stem cells (hASCs, passage p<=6) were obtained from the Radboud Biobank and cultured in Minimum Essential Medium Eagle (α-MEM) (Invitrogen, Thermo Fisher, USA). All media were supplemented with 10% fetal bovine serum (Sigma-Aldrich, USA) and 1% penicillin/streptomycin (final concentration of 100 IU mL^−1^ penicillin and 100 μg mL^−1^ streptomycin, Gibco, Thermo Fisher, USA). Dry PIC polymers were sterilized by UV for 10 min and then dissolved in the medium for 24 h at 4 °C. Cells were harvested by trypsin treatment once they reached 100% confluence and were resuspended in fresh medium. After counting, cells were mixed with the polymer solution on ice in a pre-determined ratio to achieve the required cell density and polymer concentration. After mixing, the solutions were transferred to 48-well plates (Corning, USA), 8-well chambered cover slides (Sigma-Aldrich, USA) or [μ-Slide Angiogenesis](https://ibidi.com/chambered-coverslips/41--slide-angiogenesis.html#/surface_modification-ibitreat_15_polymer_coverslip_tissue_culture_treated_sterilized/pcs_box-15) (IBIDI, Germany), and heated to 37 °C. After gel formation, warm culture medium was added onto the samples. Then all samples were subject to standard cell culture conditions (humidified atmosphere, 37 °C, 5% CO_2_).

### Matrigel experiments

For the Matrigel experiment, Corning® Matrigel® Growth Factor Reduced (GFR) Basement Membrane Matrix (Phenol red-free, LDEV-free) was used. Briefly, the Matrigel solution and cell solution were mixed in a 7:3 volume ratio on ice. The samples were incubated at 37 °C for 30 minutes before adding supernatant medium (α-MEM) on top.

### LIVE/DEAD assay

The staining protocol was adapted from the product manual (Invitrogen, Thermo Fisher, USA). In brief, first, cell culture medium on top of the gels was gently removed. Next, a 2 μM calcein AM and 4 μM EthD-1 working solution was prepared in warm α-MEM medium and added onto gel samples. After a 30-minute incubation in the cell incubator, all samples were washed by warm phosphate-buffed saline (PBS) and imaged by an Olympus FV-1000 confocal fluorescence microscope.

### Cytoskeleton staining

Gels with encapsulated cells were washed with PBS and then fixed with 4% paraformaldehyde (PFA) in PBS for 40 min. After fixation, the samples were permeabilized with 0.1% Triton X-100 in PBS for 10 min and blocked with 1% BSA in PBS for 30 min. They were then incubated with Phalloidin Atto-520 (10 μM, 1:20 in 1%BSA/PBS, Sigma-Aldrich, USA) for 1 h. All procedures were performed at 37 °C. Fluorescence images were acquired on a Leica TCS SP8 X confocal microscope, using a water objective (HC PL APO 20×/0.75, Leica and Fluotar VISIR 25×/0.95, Leica). For detection, we used a field-of-view scanner (400 Hz, bidirectional) and a hybrid photomultiplier detector (HYD-SMD, Leica). As the excitation source, a supercontinuum White Light Laser (470–670 nm, pulsed, 80 MHz, NKT Photonics) was used. The phalloidin Atto-520 was excited at 520 nm and fluorescence was collected between 530 nm and 670 nm. Images were acquired over a range of 500 μm in the *z*-direction. During measurements, the temperature was kept at 37 °C. The maximum projection images were prepared by compressing all the frames into one *z* projection using the FIJI software.

### YAP Immunostaining

Samples were gently washed with warm PBS and fixed with 4 % paraformaldehyde in PBS for 1 h at 37 °C. Next, the samples were washed with warm PBS and permeabilized with 0.1% Triton X-100 in PBS for 30 min at 37 °C. Subsequently, the samples were washed with warm PBS and blocked with 1% BSA in PBS for 24 h at 37°C. Thereafter, the samples were incubated in primary antibody (YAP Antibody (63.7): sc-101199, Santa Cruz Biotechnology, USA) 1:200 in 1 % BSA/PBS for 24 h at 37 °C and washed with warm PBS, then in secondary antibody (Goat anti-Mouse IgG (H+L) Highly Cross-Adsorbed Secondary Antibody, Alexa Fluor 633, Invitrogen, USA) 1:200 in PBS for 4 h at 37 °C. All samples were counterstained with DAPI (2.5 μg mL^−1^ in PBS) and washed with warm PBS. Fluorescence images were acquired on a Leica TCS SP8 X confocal microscope, using a water objective (HC PL APO 63×/1.20, motCORR, Leica), a hybrid photomultiplier tube as the detector (HYD-SMD, Leica) and a field-of-view scanner (200 Hz, bidirectional). As excitation sources, a supercontinuum White Light Laser (470–670 nm, pulsed, 80 MHz, NKT Photonics) was used for the Alexa Fluor 633 secondary antibody label and a UV diode laser (405 nm, pulsed, 40 MHz, PicoQuant) was used for DAPI. The Alexa Fluor 633 was excited at 632 nm and fluorescence was collected between 639 nm and 773 nm. DAPI was excited at 405 nm and fluorescence was collected between 410 nm and 551 nm. Transmission light detection was done in a separate channel using a standard Photomultiplier tube (Leica). All measurements were performed in a temperature-controlled environment (37 °C).

### Live cell imaging

A Cytosmart Lux 2 inverted bright-field microscope was placed in an incubator (37 °C, humidified) to monitor the morphology of the cells in hydrogels. Cold cell-gel mixtures (100 μL) were pipetted in a 35 mm glass-bottom dish (Cellvis, #1.5) and allowed to gelate at 37°C for 15 min. Afterward, 3 mL of CO_2_-independent α-MEM culture medium (Thermo Fisher) was pipetted on top of the samples. 24-hour monitoring was performed to acquire real-time videos with a 10× air objective. Images were acquired every 10 minutes.

### Morphology analysis

FIJI was used for analysis, images were automatically preprocessed (script 1, Supplementary information) and randomly selected cell segments were measured for morphological features (script 2, Supplementary information). A median filter was applied to the images to reduce the noise in the image while least affecting the form of the imaged cells. The filter size was chosen as small as possible to retain morphological detail, but large enough to enable a good segmentation result for the cells. One representative video per condition was used for analysis. Cell number per condition *n* = 20, except: **P0** at 1 hr (*n* = 19); **P4** at 1 hr (*n* = 14), at 3 hr (*n* = 17), at 5 hr (*n* = 19) and at 24 hr (*n* = 14). One-way analysis of variance (ANOVA) with Tuckey tests was performed for comparison between different conditions using Origin 2020.

## Data availability

All data supporting the results of this study are available in the article and the Supplementary Information Files or from the corresponding author on reasonable request.

## Acknowledgments

We thank Gaston Richelle for support on peptide synthesis, Dorien Tiemessen for support on stem cell culture, Sitara Vedaraman for support on immunostaining, Rik Nuyts for support on fluorescence imaging and Paula de Almeida for support on figure design. This project has received funding from the European Union’s Horizon 2020 research and innovation programme under the Marie Skłodowska-Curie grant agreements No 642687 (KL & DB), from the Research Foundation-Flanders (FWO, projects G0A5817N and 1529418N) and from KU Leuven (C14/16/053). KL acknowledges the EMBO Short-Term Fellowship (7427) for his research stay in KU Leuven. JV acknowledges FWO for his personal grant (1186220N).

## Author information

### Author contributions

KL: polymer synthesis, rheology, cell culture, cell encapsulation, imaging and data analysis. JV: fluorescent imaging and time-lapse bright field imaging. DB: peptide synthesis. ME: development of scripts for image analysis. EO, PT, SR, and PK supervised the project. All authors contributed to the manuscript.

## References

1 Hynes, R. O. The extracellular matrix: not just pretty fibrils. Science 326, 1216–1219, doi:10.1126/science.1176009 (2009).

2 Geiger, B., Spatz, J. P. & Bershadsky, A. D. Environmental sensing through focal adhesions. Nat. Rev. Mol. Cell Biol. 10, 21–33, doi:10.1038/nrm2593 (2009).

3 Hynes, R. O. Integrins: bidirectional, allosteric signaling machines. Cell 110, 673–687, doi:10.1016/s0092-8674(02)00971-6 (2002).

4 Doyle, A. D. & Yamada, K. M. Mechanosensing via cell-matrix adhesions in 3D microenvironments. Exp. Cell Res. 343, 60–66, doi:10.1016/j.yexcr.2015.10.033 (2016).

5 Docheva, D., Popov, C., Mutschler, W. & Schieker, M. Human mesenchymal stem cells in contact with their environment: surface characteristics and the integrin system. Journal of Cellular and Molecular Medicine 11, 21–38, doi:10.1111/j.1582-4934.2007.00001.x (2007).

6 Giancotti, F. G. & Ruoslahti, E. Integrin signaling. Science 285, 1028–1032, doi:10.1126/science.285.5430.1028 (1999).

7 Goessler, U. R. et al. Integrin expression in stem cells from bone marrow and adipose tissue during chondrogenic differentiation. Int. J. Mol. Med. 21, 271–279 (2008).

8 Desgrosellier, J. S. & Cheresh, D. A. Integrins in cancer: biological implications and therapeutic opportunities. Nature Reviews Cancer 10, 9–22, doi:10.1038/nrc2748 (2010).

9 Pierschbacher, M. D. & Ruoslahti, E. Cell Attachment Activity of Fibronectin Can Be Duplicated by Small Synthetic Fragments of the Molecule. Nature 309, 30–33, doi:DOI 10.1038/309030a0 (1984).

10 Huettner, N., Dargaville, T. R. & Forget, A. Discovering Cell-Adhesion Peptides in Tissue Engineering: Beyond RGD. Trends Biotechnol. 36, 372–383, doi:10.1016/j.tibtech.2018.01.008 (2018).

11 Bellis, S. L. Advantages of RGD peptides for directing cell association with biomaterials. Biomaterials 32, 4205–4210, doi:10.1016/j.biomaterials.2011.02.029 (2011).

12 Fong, E. & Tirrell, D. A. Collective cell migration on artificial extracellular matrix proteins containing full-length fibronectin domains. Adv. Mater. 22, 5271–5275, doi:10.1002/adma.201002448 (2010).

13 Kapp, T. G. et al. A Comprehensive Evaluation of the Activity and Selectivity Profile of Ligands for RGD-binding Integrins. Sci. Rep. 7, 39805, doi:10.1038/srep39805 (2017).

14 Bernhagen, D. et al. High-Affinity alpha(5)beta(1)-Integrin-Selective Bicyclic RGD Peptides Identified via Screening of Designed Random Libraries. Acs Comb Sci 21, 598–607, doi:10.1021/acscombsci.9b00081 (2019).

15 Kaufmann, D. et al. Chemical conjugation of linear and cyclic RGD moieties to a recombinant elastin-mimetic polypeptide--a versatile approach towards bioactive protein hydrogels. Macromol. Biosci. 8, 577–588, doi:10.1002/mabi.200700234 (2008).

16 Cipriani, F. et al. Bicyclic RGD peptides with high integrin alpha v beta 3 and alpha 5 beta 1 affinity promote cell adhesion on elastin-like recombinamers. Biomed. Mater. 14, 035009, doi:10.1088/1748-605X/aafd83 (2019).

17 Karimi, F., O’Connor, A. J., Qiao, G. G. & Heath, D. E. Integrin Clustering Matters: A Review of Biomaterials Functionalized with Multivalent Integrin-Binding Ligands to Improve Cell Adhesion, Migration, Differentiation, Angiogenesis, and Biomedical Device Integration. Adv Healthc Mater 7, e1701324, doi:10.1002/adhm.201701324 (2018).

18 Kouwer, P. H. et al. Responsive biomimetic networks from polyisocyanopeptide hydrogels. Nature 493, 651–655, doi:10.1038/nature11839 (2013).

19 Liu, K., Mihaila, S. M., Rowan, A., Oosterwijk, E. & Kouwer, P. H. J. Synthetic Extracellular Matrices with Nonlinear Elasticity Regulate Cellular Organization. Biomacromolecules 20, 826–834, doi:10.1021/acs.biomac.8b01445 (2019).

20 Ye, S. et al. A Chemically Defined Hydrogel for Human Liver Organoid Culture. Adv. Funct. Mater., 2000893, doi:10.1002/adfm.202000893 (2020).

21 Zhang, Y. et al. Polyisocyanide hydrogels as a tunable platform for mammary gland organoid formation. Adv. Sci., in press (2020).

22 op ‘t Veld, R. C. et al. Thermosensitive biomimetic polyisocyanopeptide hydrogels may facilitate wound repair. Biomaterials 181, 392–401, doi:10.1016/j.biomaterials.2018.07.038 (2018).

23 Bernhagen, D. et al. Bicyclic RGD Peptides with Exquisite Selectivity for the Integrin alpha(v)beta(3) Receptor Using a “Random Design” Approach. ACS Comb. Sci. 21, 198–206, doi:10.1021/acscombsci.8b00144 (2019).

24 Kimura, R. H., Levin, A. M., Cochran, F. V. & Cochran, J. R. Engineered cystine knot peptides that bind alpha(v)beta(3), alpha(v)beta(5), and alpha(5)beta(1) integrins with low-nanomolar affinity. Proteins-Structure Function and Bioinformatics 77, 359–369, doi:10.1002/prot.22441 (2009).

25 Schoenmakers, D. C., Rowan, A. E. & Kouwer, P. H. J. Crosslinking of fibrous hydrogels. Nat. Commun. 9, 2172, doi:10.1038/s41467-018-04508-x (2018).

26 Debets, M. F. et al. Bioconjugation with Strained Alkenes and Alkynes. Acc. Chem. Res. 44, 805–815, doi:10.1021/ar200059z (2011).

27 Strain-promoted azide-alkyne cycloaddition (SPAAC) has been proven to be simple and efficient and in our lab, a high conjugation rate is routinely realized. For this work, we did not examine the conversion of our peptide conjugations. Nevertheless, we comment here that a rapid and quantitative method is urgently needed to determine the degree of biofunctionalization at low concentrations.

28 Chaudhuri, O. et al. Hydrogels with tunable stress relaxation regulate stem cell fate and activity. Nat. Mater. 15, 326–334, doi:10.1038/Nmat4489 (2016).

29 Lam, J., Truong, N. F. & Segura, T. Design of cell-matrix interactions in hyaluronic acid hydrogel scaffolds. Acta Biomater. 10, 1571–1580, doi:10.1016/j.actbio.2013.07.025 (2014).

30 Sridhar, B. V. et al. Development of a Cellularly Degradable PEG Hydrogel to Promote Articular Cartilage Extracellular Matrix Deposition. Adv. Healthc. Mater. 4, 702–713, doi:10.1002/adhm.201400695 (2015).

31 Glorevski, N. et al. Designer matrices for intestinal stem cell and organoid culture. Nature 539, 560–564, doi:10.1038/nature20168 (2016).

32 Lam, M. T. & Longaker, M. T. Comparison of several attachment methods for human iPS, embryonic and adipose-derived stem cells for tissue engineering. J. Tissue Eng. Regen. Med. 6, s80–s86, doi:10.1002/term.1499 (2012).

33 Isomursu, A., Lerche, M., Taskinen, M. E., Ivaska, J. & Peuhu, E. Integrin signaling and mechanotransduction in regulation of somatic stem cells. Exp. Cell Res. 378, 217–225, doi:10.1016/j.yexcr.2019.01.027 (2019).

34 Jokinen, J. et al. Integrin-mediated cell adhesion to type I collagen fibrils. J. Biol. Chem. 279, 31956–31963, doi:10.1074/jbc.M401409200 (2004).

35 Hynes, R. O. Integrins: versatility, modulation, and signaling in cell adhesion. Cell 69, 11–25, doi:10.1016/0092-8674(92)90115-s (1992).

36 Dupont, S. et al. Role of YAP/TAZ in mechanotransduction. Nature 474, 179–183, doi:10.1038/nature10137 (2011).

37 Lee, J. Y. et al. YAP-independent mechanotransduction drives breast cancer progression. Nat. Commun. 10, 1–9, doi:10.1038/s41467-019-09755-0 (2019).

38 de Lucas, B., Perez, L. M. & Galvez, B. G. Importance and regulation of adult stem cell migration. J Cell Mol Med 22, 746–754, doi:10.1111/jcmm.13422 (2018).

39 van Helvert, S., Storm, C. & Friedl, P. Mechanoreciprocity in cell migration. Nat. Cell Biol. 20, 8–20, doi:10.1038/s41556-017-0012-0 (2018).

40 Wisdom, K. M. et al. Matrix mechanical plasticity regulates cancer cell migration through confining microenvironments. Nat. Commun. 9, 1–13, doi:10.1038/s41467-018-06641-z (2018).

41 Jacquemet, G., Hamidi, H. & Ivaska, J. Filopodia in cell adhesion, 3D migration and cancer cell invasion. Curr. Opin. Cell Biol. 36, 23–31, doi:10.1016/j.ceb.2015.06.007 (2015).

42 Petrie, R. J. & Yamada, K. M. At the leading edge of three-dimensional cell migration. J. Cell Sci. 125, 5917–5926, doi:10.1242/jcs.093732 (2012).

43 Veevers-Lowe, J., Ball, S. G., Shuttleworth, A. & Kielty, C. M. Mesenchymal stem cell migration is regulated by fibronectin through alpha(5)beta(1)-integrin-mediated activation of PDGFR-beta and potentiation of growth factor signals. J. Cell Sci. 124, 1288–1300, doi:10.1242/jcs.076935 (2011).

