## Supplementary Information for "Rapid stem cell spreading induced by high affinity α_5_β_1_ integrin-selective bicyclic RGD peptide in biomimetic hydrogels"

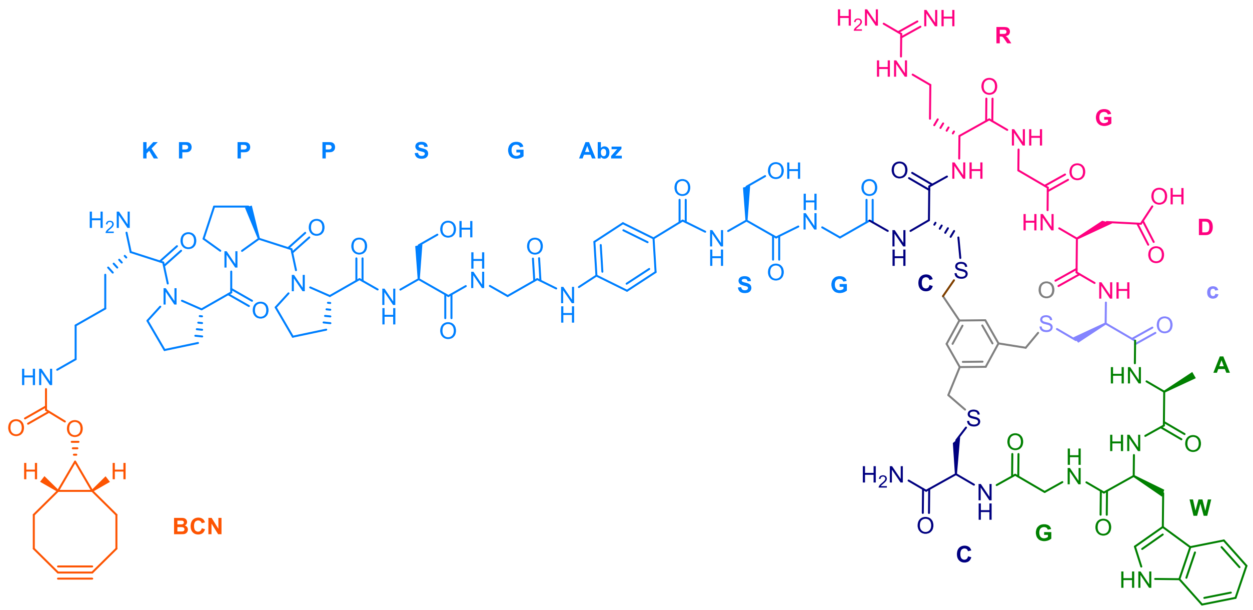


**Supplementary Figure S1. Molecular structure of the bicyclic peptide-BCN construct.** As an example, the peptide in gel **P8** is represented. The color follows Figure 1: BCN in orange; spacer in blue; bicycle-scaffold gray; *L*-cysteine (bark blue) and *D*-cysteine (violet) that link the peptide chain to the scaffold; RGD binding domain in pink; second peptide sequence in green.

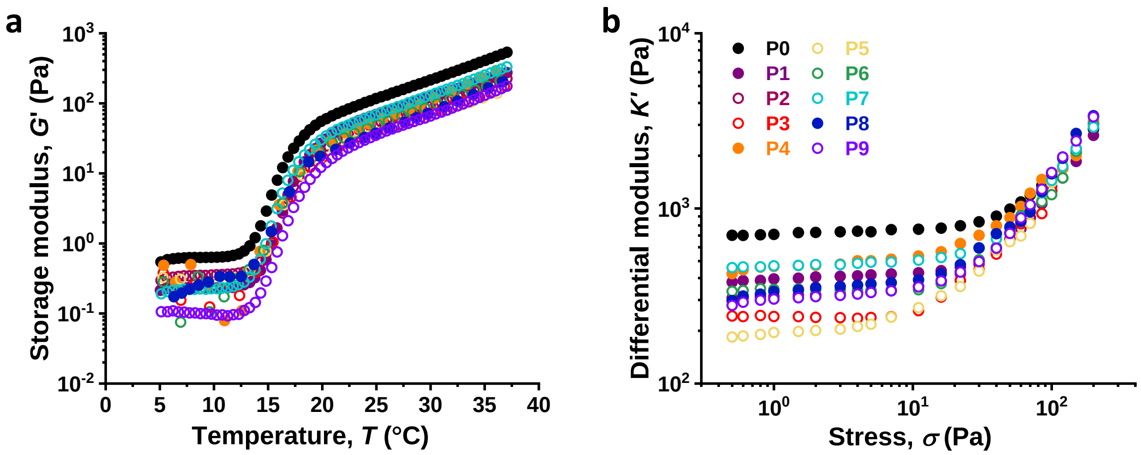


**Supplementary Figure S2. Mechanical characterization of gels. a.** Temperature ramps show that all gels are formed upon a similar lower critical solution temperature (LCST) at ~15 °C. **b.** Differential modulus as a function of the externally applied stress. Data suggests that all gels maintain the intrinsic nonlinear mechanics regardless of the conjugated peptides.

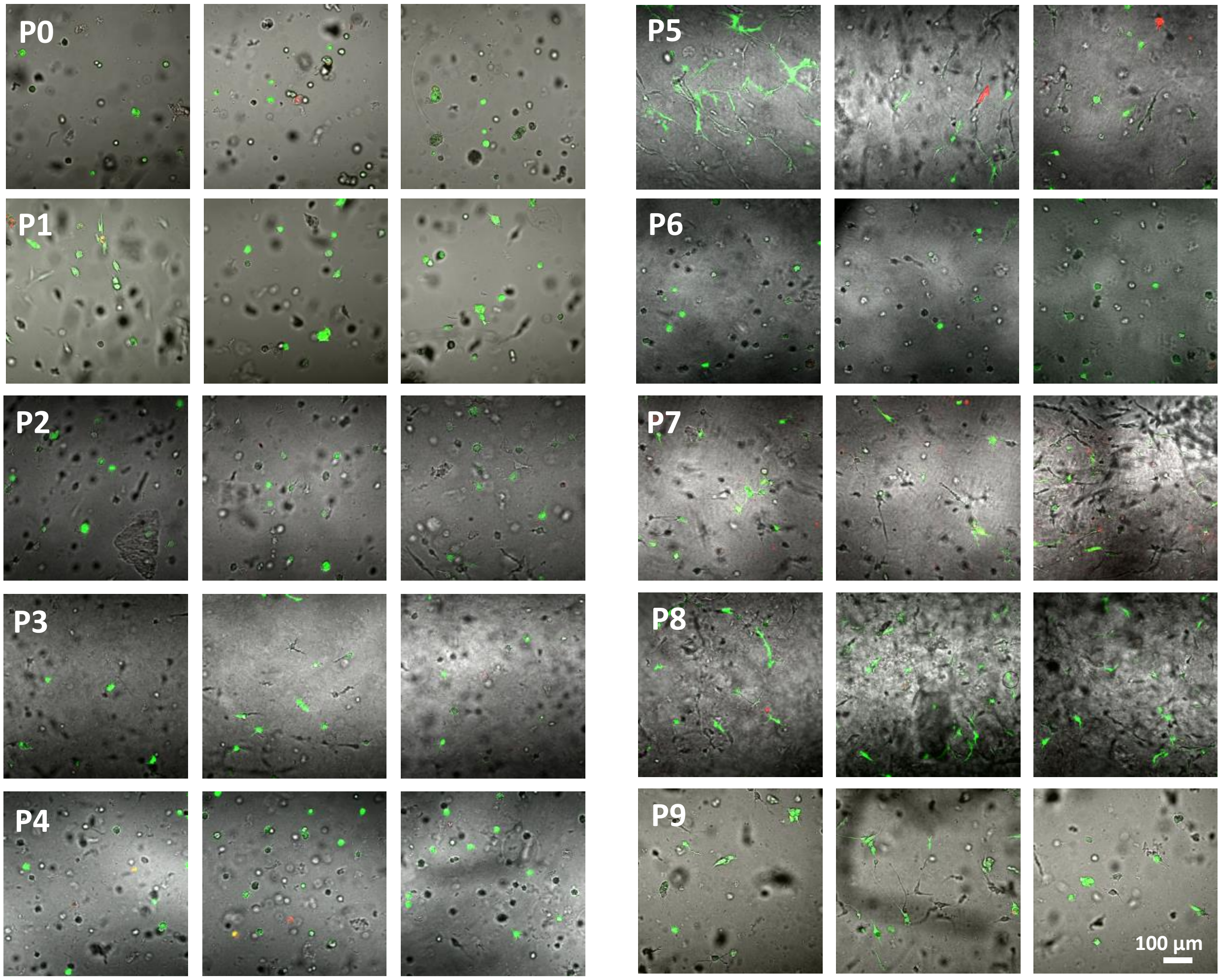


**Supplementary Figure S3. LIVE/DEAD staining of stem cells 72 h after encapsulation in P0–P9.** 3 representative slides at different z levels are shown (green: live; red: dead). All samples show good cell viability.

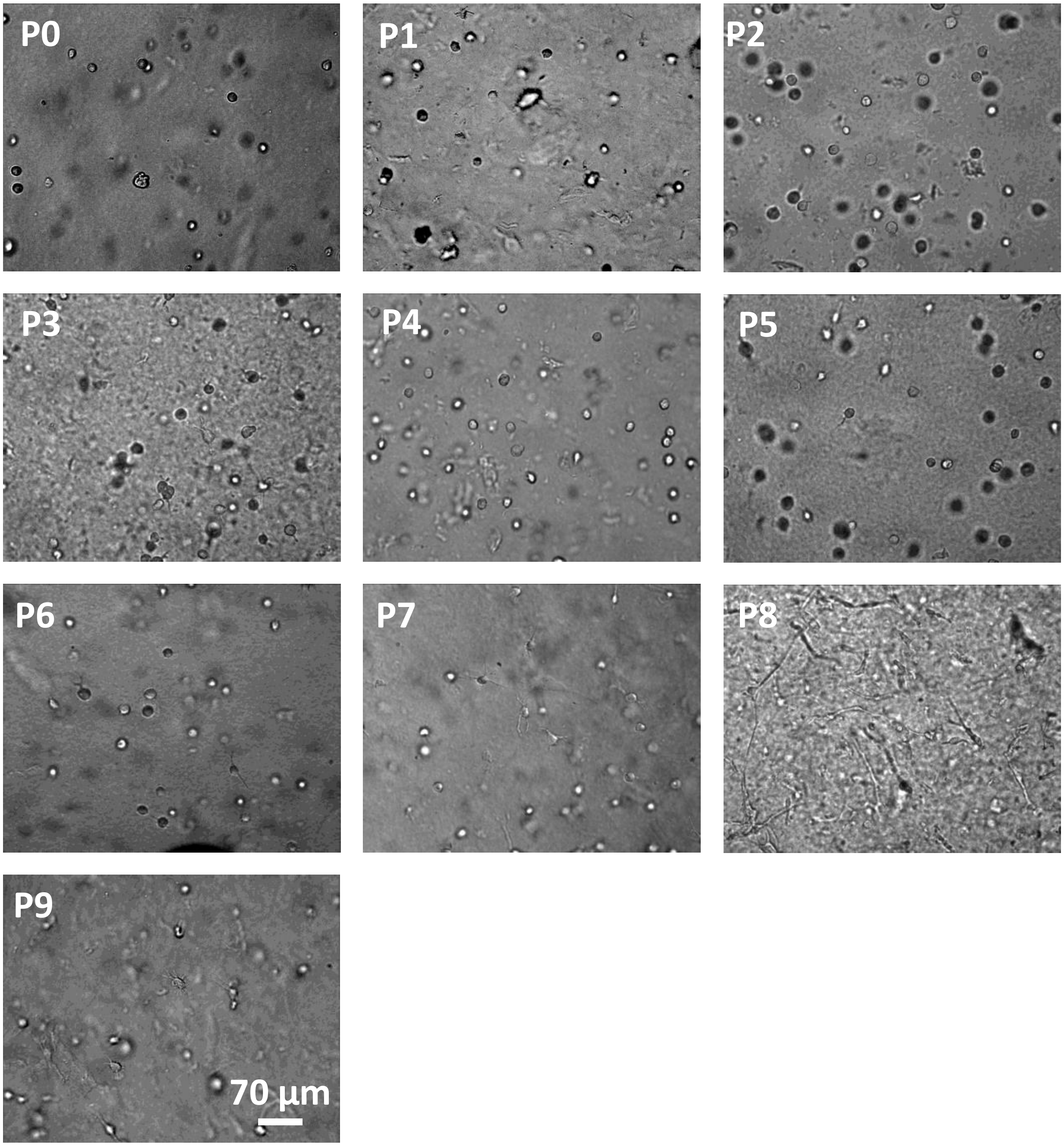


**Supplementary Figure S4. Bright field images of human adipose-derived stem cells (hASCs) 24 h after encapsulation in P0–P9.** Cells in **P8** show a significant morphological change.

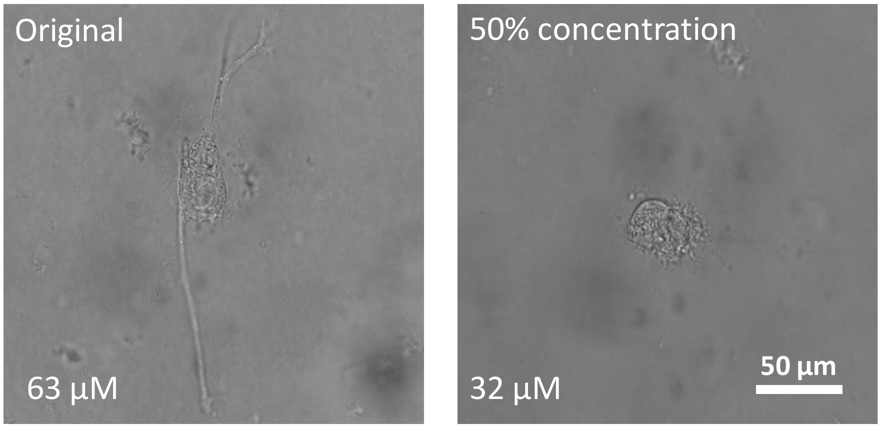


**Supplementary Figure S5. Influence of peptide density on cell spreading.** The rapid cell spreading 24 h after cell seeding disappear when the peptide density is halved from 63 to 32 μM.

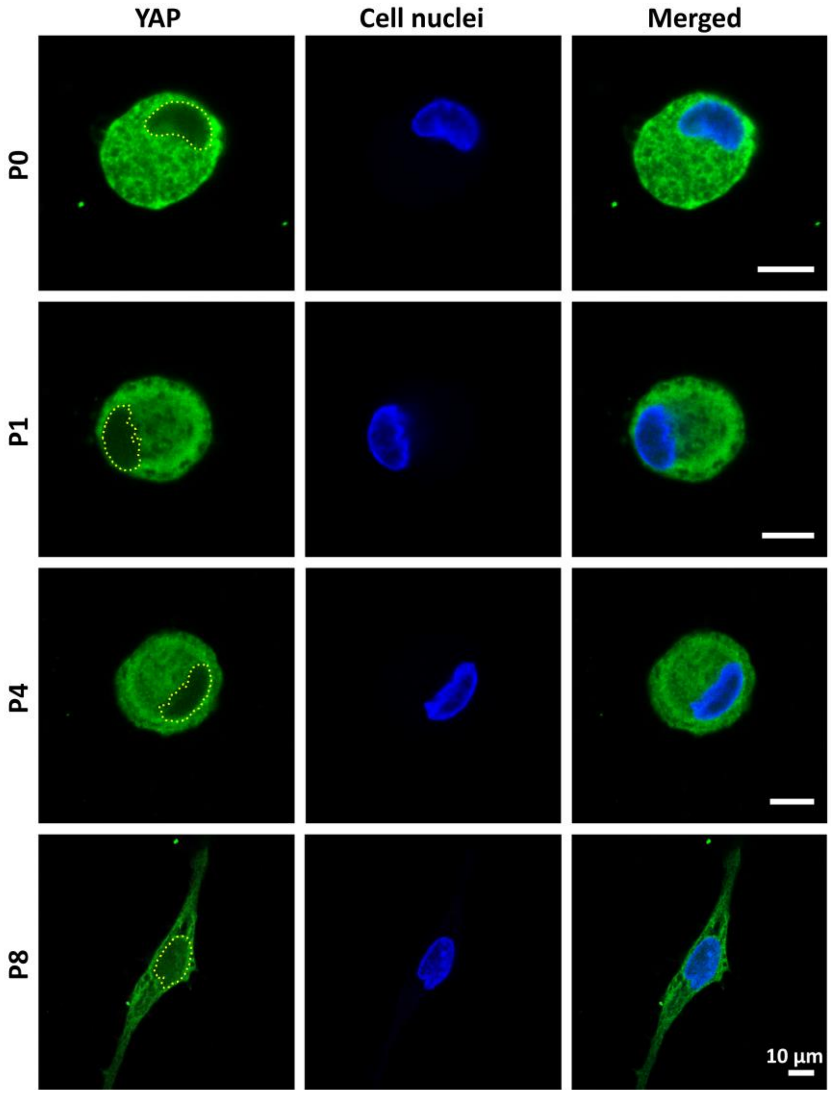


**Supplementary Figure S6. YAP staining 24 h after encapsulation in P0, P1, P4, and P8.** Nuclear localization of YAP is not seen in any of the matrices.

### Further Supplementary Material

**Supplementary Movies M1 – M5. Time lapse bright field microscopy.** First 24 hours of the 3D cultures of hACSs (200,000 cells $mL^{-1}$) in **P0**, **P1**, **P4**, **P8** and Matrigel.

***Code of Script 1:***

```
roiManager("Reset");
run("Clear Results");
run("Duplicate...", "title=ChosenSlice");
run("Invert");
run("Median...", "radius=4");
setAutoThreshold("Moments dark");
setOption("BlackBackground", false);
run("Convert to Mask");
run("Fill Holes", "slice");
run("Analyze Particles...", "display add");
roiManager("Show None");
```

***Code of Script 2:***

```
Roi.getCoordinates(xpoints, ypoints);
nrOfPoints = xpoints.length;
chosenRois = newArray(nrOfPoints);
nrOfRois = roiManager("count");
pickedRois = 0;
for(roiNr = 0; roiNr < nrOfRois; roiNr++)
{
	pointNotFound = true;
	roi = roiManager("select", roiNr);
	for(pointNr = 0; (pointNr < nrOfPoints) || !pointNotFound ; pointNr++)
	{
		if(Roi.contains(xpoints[pointNr], ypoints[pointNr]))
		{
			chosenRois[pickedRois] = roiNr;
			pickedRois++;
			pointFound = false;
		}
	}
}
Table.create("Measurements");
roiManager("Select", chosenRois);
roiManager("Combine");
for(roiNr = 0; roiNr < nrOfPoints; roiNr++)
{
	picked = chosenRois[roiNr];
	Table.set("Roi Number", roiNr, picked);
	area = getResult("Area", picked);
	Table.set("Area", roiNr, area);
	circ = getResult("Circ.", picked);
	Table.set("Circularity", roiNr, circ);
}
```
